# Functional connectivity scaffolds the emergence of the visual word form area

**DOI:** 10.64898/2026.08.14.744903

**Authors:** Maya Yablonski, Jamie L. Mitchell, Hannah L. Stone, Mia Jimenez, Jason D. Yeatman

## Abstract

Complex cognitive tasks, like reading, require coordinated computations across distant cortical regions. Connectivity among regions is therefore key for understanding the neurobiological basis of behavior. It remains an open question whether connectivity serves as an innate organizational blueprint that guides functional specialization, or if it evolves with learning. Here, we address this fundamental question through the prism of learning to read: We measured functional connectivity at five timepoints over a year in children with dyslexia who participated in an intensive reading intervention. We found that specific patterns of functional connectivity of the reading circuitry were in place prior to the intervention, and remained stable over time. Further, functional connectivity before the intervention predicted the location of text-selective regions a year later, showing that connectivity precedes and predicts functional specialization that emerge with learning. These findings support a view of functional connectivity as a stable organizational principle of high-level cortex.

## Introduction

A key goal of cognitive neuroscience is understanding the guiding principles of the brain’s functional organization. A fundamental question concerns the development of specialized cortical regions that are tuned to specific visual categories, like faces, places and words ^1–3^. Despite individual differences, these regions can be consistently identified in typical locations within ventral occipitotemporal cortex (VOTC). Several theories have been proposed to explain the sources of this functional mosaic in terms of the computations carried out by each area, for example their organization by eccentricity ^4–6^ or sensitivity to visual features ^7,8^. On the other hand, others proposed that the function of a region is determined primarily by its long range connections ^9,10^. According to this ‘connectivity hypothesis’, innate connectivity patterns, which form each area’s inputs and outputs, determine the optimal location for category-selective regions to emerge ^3^.

The connectivity hypothesis is supported by the surprising correspondence between functional connectivity maps and task activations ^11,12^. Recent studies have shown that functional connectivity at rest can predict individual differences in task evoked activations across a variety of tasks ^13–15^. Further, adjacent category-selective regions were shown to have distinct profiles of functional connectivity with distant cortical regions ^16–19^. Importantly, these studies have been conducted in healthy adults, predicting task activations from resting state (task-free) data collected at the same scanning session. It thus remains an open question whether the coupling between connectivity and function is an intrinsic property of the brain, or arises over development due to experience. If connectivity is innate, it may serve as a scaffold that guides the emergence of category-selective regions given sufficient experience and training. Alternatively, connectivity and function may co-develop, such that accumulating experience processing a specific category drives both the specialization of a dedicated region, and the strengthening of its connections with relevant systems throughout the cortex.

Some clues to this developmental question come from studies that observed adult-like category specific functional connectivity patterns in the brains of infants ^20–22^. These observations show that the general landscape of VOTC functional connectivity exists at birth, before functional specialization has taken place. On a similar vein, studies have found that functional connectivity in the visual cortex of the congenitally blind follows a retinotopic organization ^23^, and functional connectivity in the auditory cortex in the deaf follows a tonotopic organization ^24^, showing that typical connectivity patterns exist in the absence of sensory experience. While this supports a view of functional connectivity as an innate architecture, critical evidence is missing as to how this connectivity guides the subsequent emergence of category-selective regions, and what role experience and learning play in shaping these connections. To fill this gap, longitudinal data that specifically track functional connectivity before and after category-selectivity develops are necessary.

Learning to read provides a unique testbed for these questions. Reading is a new cultural invention and takes years of formal instruction and practice to master. Thus, the brain is not pre-wired for reading in the same sense that it may be pre-wired for processing other ‘natural’ visual categories, or spoken language ^25^. Rather, as children learn to read, they have to form new mappings between arbitrary written visual symbols, and language units (sounds and meanings). This process leads to the emergence of the visual word form area (VWFA; ^26,27^) which has been the focus of extensive research in the field. Recent work has further suggested that the VWFA comprises two distinct subregions, VWFA-1 and VWFA-2, which differ in their functional properties and anatomical connections ^28,29^. Specifically, VWFA-2 is thought to be a hub that bridges the visual system with the language system ^30,31^. In line with this idea, we recently found that VWFA-2 shows specific coupling with frontal language regions in the inferior frontal cortex (IFC) in adults and children with a wide range of reading skills ^32^. It remains unclear whether these connectivity patterns precede and scaffold the emergence and specialization of VWFA-2, or whether it’s the consequence of learning to read that elevates this specific connection beyond its neighboring areas.

To address this gap, here we leverage a longitudinal intervention study, where children with dyslexia participated in an intensive intervention program to improve their reading skills. We found that many children with dyslexia did not have a VWFA prior to the intervention, and that the intervention drives the emergence of the VWFA for some participants ^33^. The intervention settings and this unique population allow us to ask: 1) Is there a connectivity blueprint linking the visual and language networks that predates the ability to read proficiently? 2) Do these connectivity patterns change with learning? 3) Can connectivity predict the emergence of text-selectivity as it unfolds over time? If functional connectivity of the VWFA-2 is formed by learning to read, we would expect to see a difference between typical readers and children with dyslexia, and more specifically with children who haven’t developed text-selectivity (no VWFA). We would also expect to see functional connectivity change as text-selectivity emerges following the intervention. On the other hand, if connectivity is an innate feature scaffolding the development of functional modules, we would expect it to be consistent across different populations, remain stable over time, and predict the location of the VWFA in the future.

## Results

In this longitudinal study, we scanned 44 children with dyslexia who completed a targeted 8-week reading intervention program, and 43 controls with (N=19) and without dyslexia (N=24). Our previous work established the efficacy of the intervention in increasing reading scores with large effect sizes ^33^. All children (ages 7.4-13.8 years, 44/43 Male/Female) completed up to 5 functional magnetic resonance imaging (fMRI) scans before the intervention, immediately after its completion, six months and twelve months later. This dense sampling scheme allows us to delineate a trajectory of both short- and long-term changes in brain function and connectivity following the intervention. In each timepoint, children completed a functional localizer where they viewed text and non-text stimuli while performing tasks that did not explicitly require reading ^33,34^. In addition, they completed a task-free resting state scan where they watched a silent nature movie without language content. Movie watching was chosen over classical rest due to evidence that movies reduce motion, improve subject compliance, and improve prediction ability, all critical in developmental populations ^35–37^. The functional localizer was used to define text-selective regions of interest (ROIs): The subregions of the visual word form area (VWFA-1 and VWFA-2), as well as frontal language regions in the inferior frontal cortex (IFC). These ROIs were used as seeds in the functional connectivity analysis of the independent movie-watching data, to explore the functional connectivity patterns of the reading network and how they change following intervention and over the course of a year.

### VWFA-2 shows a unique connection with frontal language areas, that is agnostic to reading ability and text-selectivity

We first sought to determine whether VWFA-2 shows privileged connectivity with the IFC in children with dyslexia prior to the intervention. To this end, we created difference maps subtracting, vertex-wise, the connectivity strength of VWFA-2 from that of VWFA-1. Visual inspection of these maps in the typical reader group (N=19 after data exclusion; Figure 1) revealed the expected pattern of greater VWFA-2 connectivity in the frontal lobe, primarily around the inferior frontal sulcus (IFS) and precentral gyrus (PrG), as well as in the supramarginal gyrus (SMG), replicating our previous findings ^32^. Next, we repeated this procedure in the dyslexic group (N=53, combining both intervention and control children with dyslexia) and found similar connectivity patterns. Importantly, these difference maps were not statistically different between the typical and dyslexic groups (Welch’s independent samples t-test, no vertex survived FDR correction at a level of 0.05). The same was true when we directly compared each seed ROI’s connectivity maps between groups (for the seeds of VWFA-1, VWFA-2, and IFC). This extends previous findings, showing that the unique link between VWFA-2 and frontal language regions exists in children who are struggling with learning to read (i.e., dyslexia).

**Figure 1.**
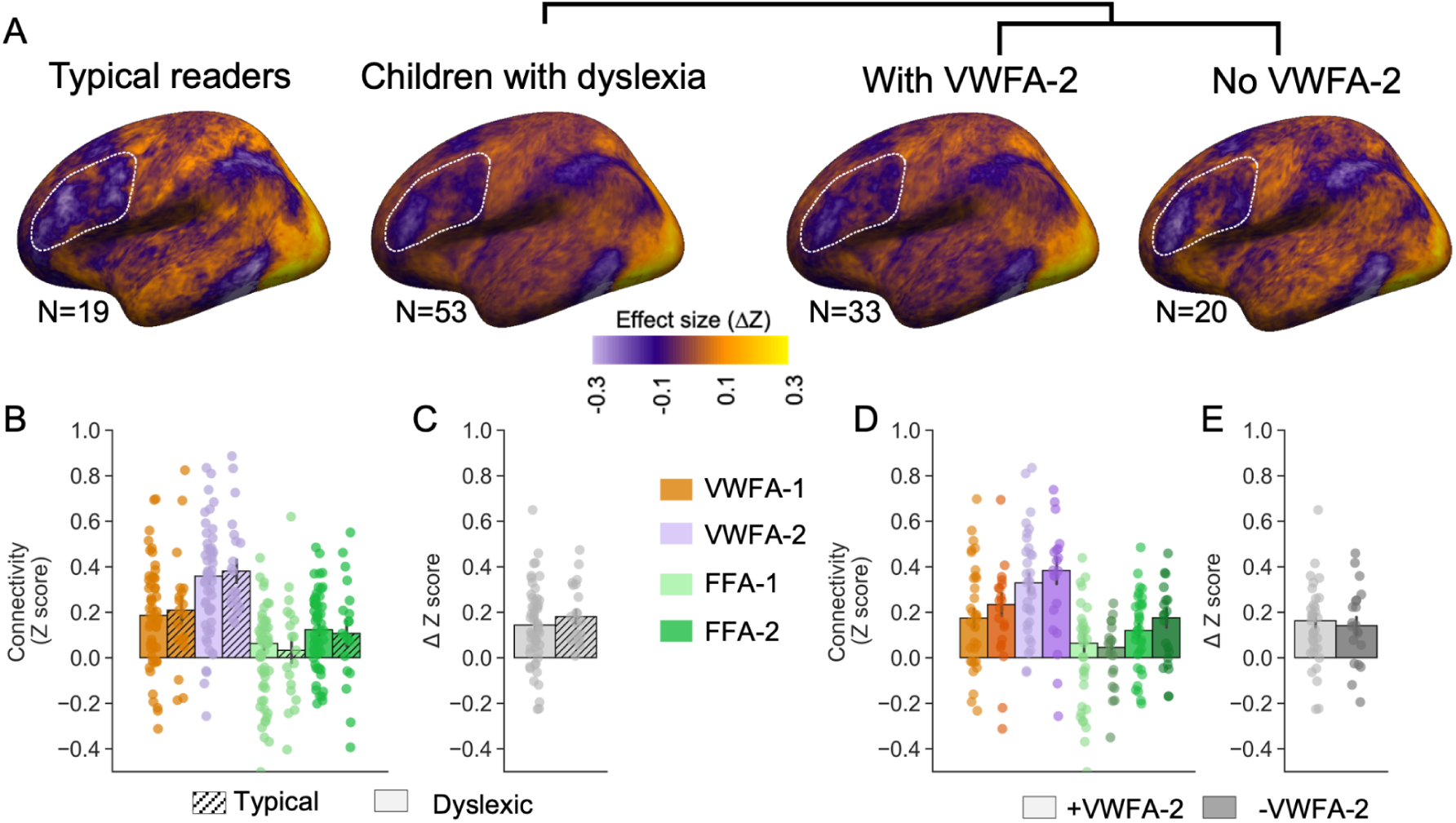
VWFA-2 is uniquely linked to the frontal lobe regardless of reading ability and text-selectivity. (A) Group difference maps comparing whole brain connectivity of VWFA-2 with whole brain connectivity of VWFA-1, for typical readers and in children with dyslexia (left), and when splitting the dyslexia group by presence of a text-selective VWFA-2 (right). Maps show the effect size in terms of connectivity difference (Δ z-score), where negative values denote greater VWFA-2 connectivity (purple) and positive values denote greater VWFA-1 connectivity (orange). White dashed contour highlights the frontal cortex. (B) ROI to ROI connectivity between the inferior frontal cortex (IFC) and each of the ventral ROIs. Solid bars represent children with dyslexia (N=53), patterned bars represent typical readers (N=19). (C) Difference in connectivity strength between IFC-VWFA-2 compared with IFC-VWFA-1 (Z(IFC-VWFA-2) – Z(IFC-VWFA-1). (D) ROI to ROI connectivity between the IFC and each of the ventral ROIs, in children who had a VWFA-2 at the pre-intervention timepoint (+VWFA2, lighter shades; N=33), and children who didn’t have a text-selective VWFA-2 pre-intervention (-VWFA-2, darker shades; N=20). (E) Difference in connectivity strength between IFC-VWFA-2 compared with IFC-VWFA-1 (Z(IFC-VWFA-2) – Z(IFC-VWFA-1). Throughout the figure, bar height denotes the median and error bars denote the standard error of the mean.

To directly quantify connectivity strength between the IFC and the ventral regions, we examined ROI-to-ROI connectivity using linear mixed effect (LME) models comparing connectivity strength between IFC and VWFA-2, versus all other ventral ROIs, with an interaction term for reading group (with/without dyslexia). This revealed again a main effect of ROI, such that IFC connectivity was strongest with VWFA-2 (VWFA-1: *β* = −0.14, p < 0.001; FFA-1: *β* = −0.32, p < 0.001, FFA-2: *β* = −0.21, p < 0.001), while the group effect was not significant (*β* = 0.08, p = 0.175), nor was the interaction between group and ROI (see Table 1; Figure 1, bottom panel). To confirm, we also ran separate models within each group (typical readers, dyslexic readers) and found in each of them the same effects: IFC had stronger connectivity with VWFA-2 compared with the other ventral ROIs, in the typical reading group (see Supplementary Table S1; VWFA-1: *β* = −0.20, p < 0.0001; FFA-1: *β* = −0.38, p < 0.0001; FFA-2: *β* = −0.30, p < 0.0001) as well as in the dyslexic group (VWFA-1: *β* = −0.14, p < 0.0001; FFA-1: *β* = −0.32, p < 0.0001; FFA-2: *β* = −0.21, p < 0.0001). Lastly, we calculated for each child the difference between their IFC-VWFA-1 connectivity and their IFC-VWFA-2 connectivity (*Δ*IFC). An independent samples t-test between the typical and dyslexic groups revealed that this connectivity difference was similar regardless of reading ability (t_(df=41.98)_=1.542, p = 0.131; Figure 1). Together this confirms that pre-intervention, the link between IFC and VWFA-2 is greater than the neighboring ROIs, and this is not affected by reading ability.

**Table 1.**
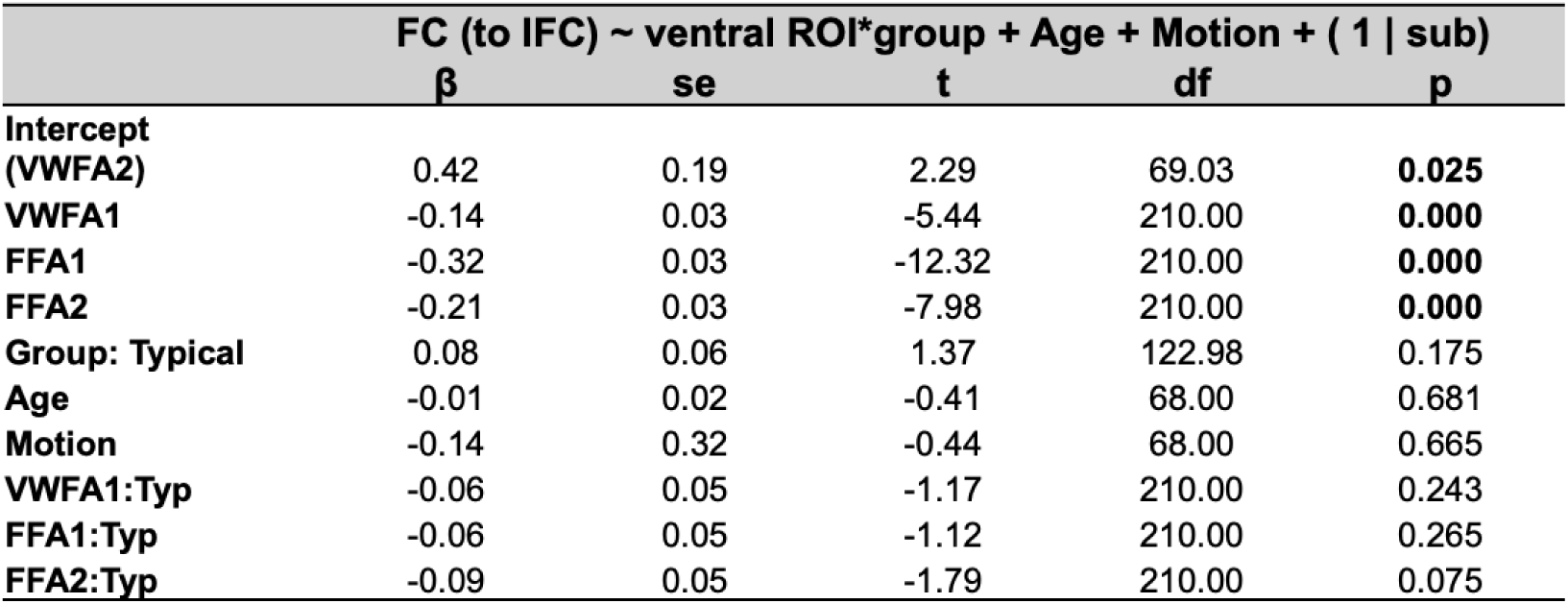
Linear mixed effect models in the pre-intervention timepoint, comparing functional connectivity between each of the ventral ROIs (VWFA-1, VWFA-2, FFA-1, FFA-2) and the inferior frontal cortex (IFC), controlling for reading group. The model was run on all children with available data at the pre-intervention timepoint (N=72). Age- number of years at the pre-intervention visit. Motion- mean framewise displacement (mm). VWFA-2 was set as the reference level. Data were extracted from template ROIs.

We next aimed to investigate whether the unique relationship between the VWFA-2 area and the frontal lobe depends on text selectivity. To this end, we focused on a specific subgroup of children with dyslexia who did not have a discernible text-selective VWFA-2 prior to the intervention. We used an independent category localizer to identify in each individual child their text-selective regions (VWFA-1, VWFA-2) and face-selective regions (FFA-1, FFA-2) based on their responses to visually presented stimuli. Using this method, we identified a subgroup of children with dyslexia who did not show text-selective responses in VOTC ^33^. We used this unique sample to investigate whether the cortical area where VWFA-2 is typically found (when defined using templates or atlases) shows greater connectivity with IFC even in the absence of text-selectivity. We thus split the dyslexic group to children with a text-selective VWFA-2 at the pre-intervention timepoint (*+VWFA2*, N=33), and children whose VWFA-2 could not be identified based on text-selectivity at that time (*-VWFA2*, N=20). The difference maps subtracting VWFA-2 from VWFA-1 connectivity were strikingly similar between these two groups (Figure 1, bottom panel). In fact, the difference maps were not statistically different between these groups and nor were the connectivity maps for each of the seed ROIs (VWFA-1, VWFA-2, IFC).

To directly quantify the effect of having a text-selective VWFA-2, we ran a LME model where we evaluated the functional connectivity between IFC and the VWFA-2 area compared with all other ROIs, with an interaction term for VWFA-2 presence. This revealed again a main effect of ROI, such that IFC connectivity was strongest with VWFA-2 (VWFA-1: *β* = −0.14, p = 0.002; FFA1: *β* = −0.32, p < 0.001, FFA-2: *β* = −0.20, p < 0.001), while the effect of VWFA-2 presence (whether VWFA-2 could be identified based on text-selectivity) was not significant (*β* = 0.00, p = 0.993), nor was the interaction between VWFA-2 presence and ROI (see Table 2; Figure 1, bottom panel). To confirm, we also ran separate models within each group (*+VWFA2* and *-VWFA2*) and found in each of them the same effects (Supplementary Table S2). Lastly, we calculated for each child the difference between their IFC-VWFA1 connectivity and their IFC-VWFA2 connectivity (*Δ*IFC). An independent sample t-test between the *+VWFA2* and *-VWFA2* groups revealed that this connectivity difference was similar regardless of VWFA-2 text- selectivity (t_(df=41.55)_ = 0.0361, p = 0.9713; Figure 1). Together, our findings show that VWFA-2 has a unique relationship with IFC, regardless of reading ability and text-selectivity.

**Table 2.**
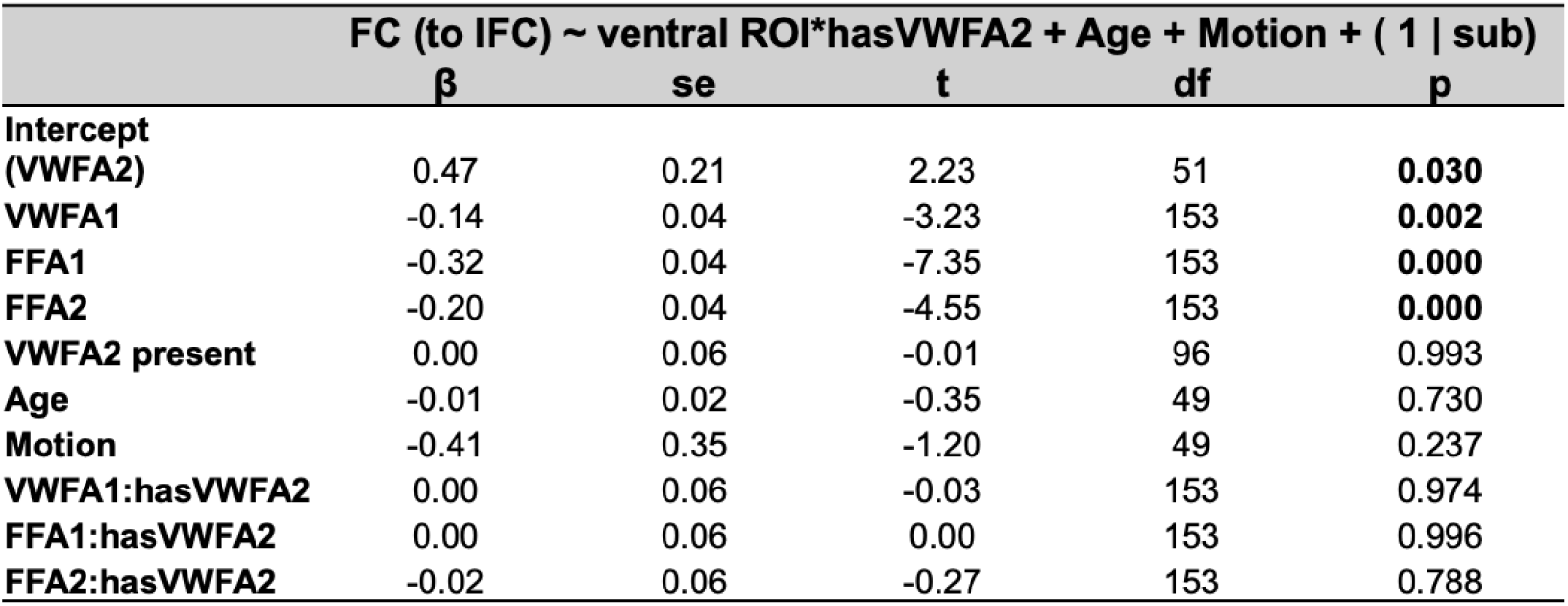
Linear mixed effect models in the pre-intervention timepoint, comparing functional connectivity between each of the ventral ROIs (VWFA-1, VWFA-2, FFA-1, FFA-2) and the inferior frontal cortex (IFC), controlling for VWFA2 presence. The model was run on all children with dyslexia at the pre-intervention timepoint (N=53). Age- number of years at the pre-intervention visit. Motion- mean framewise displacement (mm). VWFA-2 was set as the reference level. Data were extracted from template ROIs.

### Elevated connectivity between VWFA-2 and IFC is stable over time

We next examined whether IFC-VWFA-2 connectivity changes following the intervention. Focusing on the intervention group alone pre-intervention (N=37), linear mixed effects models controlling for the effects of age and motion confirmed again that the IFC had stronger connectivity with VWFA-2 compared with all other ventral ROIs tested (VWFA-1: *β* = −0.13, p = 0.0004; FFA-1: *β* = −0.31, p < 0.0001; FFA-2: *β* = −0.22, p < 0.0001). This was also the case in all other timepoints (see Supplementary Table S3, Figure 2).

**Figure 2.**
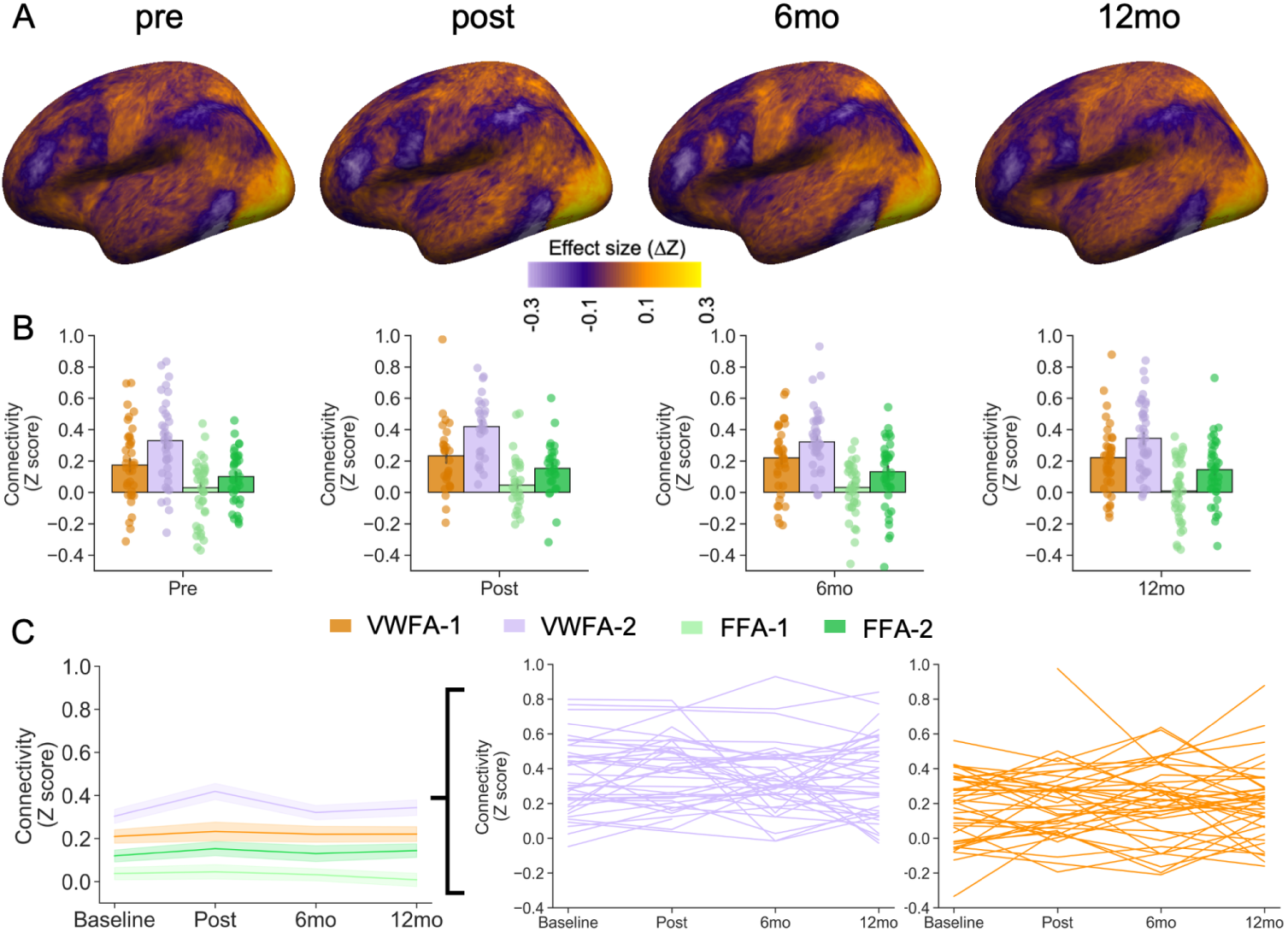
Connectivity profiles are stable with learning. (A) Group maps (N=37; N=28; N=36; N=39) comparing whole brain connectivity of VWFA-2 with whole brain connectivity of VWFA-1 in children with dyslexia who completed an intensive reading intervention, shown across the study timepoints. Maps show the effect size in terms of connectivity difference (Δ z-score), where negative values denote greater VWFA-2 connectivity (purple) and positive values denote greater VWFA-1 connectivity (orange) (B) ROI-to-ROI connectivity between the text-selective inferior frontal cortex (IFC) and the ventral ROIs. Bar height denotes the median values while each point represents an individual subject. In all timepoints, VWFA2-IFC connectivity (lilac bar) is significantly greater than the IFC connectivity with the rest of the ventral ROIs (see Supplementary Table S3). (C) Connectivity with the IFC over time. Shown are group averages (left) and individual trajectories (right). There was no significant change in connectivity over time (see Table 3).

**Table 3.** Longitudinal linear mixed effect models in the intervention group (N=44). The top model treats time as a continuous variable to examine linear time effects. The bottom models treat time as a categorical variable, comparing each session to the baseline. No significant time effects were observed in either model affecting functional connectivity of the VWFAs and the IFC. Age- number of years at the pre-intervention visit. Motion- mean framewise displacement (mm).

| FC (VWFA1-IFC) ~ Time (weeks) + Age + Motion + (1 \| sub) |  |  |  |  |  | FC (VWFA2-IFC) ~ Time (weeks) + Age + Motion + (1 \| sub) |  |  |  |  |  |
| --- | --- | --- | --- | --- | --- | --- | --- | --- | --- | --- | --- |
| | $\beta$ | se | t | df | p | | $\beta$ | se | t | df | p |
| (Intercept) | 0.04 | 0.26 | 0.15 | 43.64 | 0.8822 |  | 0.19 | 0.27 | 0.72 | 42.95 | 0.4773 |
| Time (weeks) | 0.00 | 0.00 | 0.61 | 138.99 | 0.5458 |  | 0.00 | 0.00 | -1.17 | 138.31 | 0.2423 |
| Age | 0.01 | 0.03 | 0.47 | 40.32 | 0.6380 |  | 0.02 | 0.03 | 0.66 | 39.78 | 0.5114 |
| Motion | 0.18 | 0.26 | 0.68 | 173.81 | 0.4994 |  | -0.01 | 0.26 | -0.04 | 173.06 | 0.9717 |

**Table 3.** Longitudinal linear mixed effect models in the intervention group (N=44).
| FC (VWFA1-IFC) ~ Time (session) + Age + Motion + (1 \| sub) |  |  |  |  |  | FC (VWFA2-IFC) ~ Time (session) + Age + Motion + (1 \| sub) |  |  |  |  |  |
| --- | --- | --- | --- | --- | --- | --- | --- | --- | --- | --- | --- |
| | $\beta$ | se | t | df | p | | $\beta$ | se | t | df | p |
| (Intercept) | 0.02 | 0.26 | 0.09 | 44.20 | 0.9280 |  | 0.16 | 0.27 | 0.59 | 43.41 | 0.5585 |
| session: post | 0.03 | 0.04 | 0.85 | 137.55 | 0.3950 |  | 0.06 | 0.04 | 1.65 | 136.70 | 0.1006 |
| session: 6mo | 0.01 | 0.03 | 0.44 | 137.32 | 0.6573 |  | -0.02 | 0.03 | -0.63 | 136.54 | 0.5309 |
| session: 12mo | 0.02 | 0.03 | 0.71 | 136.68 | 0.4789 |  | -0.01 | 0.03 | -0.39 | 135.95 | 0.7004 |
| Age | 0.01 | 0.03 | 0.50 | 40.45 | 0.6199 |  | 0.02 | 0.03 | 0.72 | 39.91 | 0.4736 |
| Motion | 0.19 | 0.26 | 0.72 | 171.85 | 0.4725 |  | 0.01 | 0.26 | 0.06 | 170.75 | 0.9544 |

After establishing the pattern cross-sectionally within each timepoint, we next examined whether functional connectivity strength between IFC and the VWFAs change with time within the intervention group. To this end, we ran LMEs predicting functional connectivity strength between IFC and either VWFA-1 or VWFA-2 as a function of time, controlling for age and motion. There was no significant time effect when time was treated as a continuous variable (i.e., number of weeks since the pre-intervention visit, VWFA-1: *β* = 0.00, p = 0.5458; VWFA-2: *β* = 0.00, p = 0.2423; Table 3). To account for non-linear effects of time, we repeated these analyses while treating timepoint as a categorical variable, such that each visit was compared to the baseline, coded as the average of the two pre-intervention visits. Again, the effect of timepoint was not significant for either VWFA-1(*β* = 0.03, p = 0.3950; *β* = 0.01, p = 0.6573; *β* = 0.02, p = 0.4789, for post-intervention, 6 month and 12 month followup, respectively) or VWFA-2 (*β* = 0.06, p = 0.1006; *β* = −0.02, p = 0.5309; *β* = −0.01, p = 0.7004, for post-intervention, 6 month and 12 month followup, respectively). In sum, we did not find evidence for change in functional connectivity between the VWFAs and IFC following reading intervention.

Due to accumulating evidence that the location of the VWFA is highly variable between individuals ^38,39^, a potential concern could be that the template ROIs are not sensitive enough to capture learning related changes in functional connectivity. To address this possibility, we repeated the above analyses using the individual ROIs we defined based on text selectivity (see *Methods* for details about region of interest definition). This analysis revealed similar results: Cross sectionally, IFC had stronger connectivity with VWFA-2 compared with all other ventral ROIs tested before the intervention (VWFA-1: *β* = −0.18, p = 0.0001; FFA-1: *β* = −0.38, p < 0.0001; FFA-2: *β* = −0.32, p < 0.0001). This was also the case in all other timepoints (see Supplementary Table S4). We then turned to longitudinal models, where we did not observe a significant effect of time on functional connectivity (time continuous: VWFA-1: *β* = 0.00, p = 0.2718; VWFA-2: *β* = 0.00, p = 0.1075; Supplementary Table S5). The same was true when time was treated as a categorical variable (VWFA-1: *β* = 0.03, p = 0.5414; *β* = 0.03, p = 0.3909; *β* = 0.03, p = 0.4097, for post-intervention, 6 month and 12 month followup, respectively; VWFA-2: *β* = 0.02, p = 0.7001; *β* = −0.02, p = 0.5706; *β* = −0.07, p = 0.0945, for post-intervention, 6 month and 12 month followup, respectively). Together, this shows that even when individually defining text-selective ROIs, the functional connectivity between them remains stable.

### Intervention drives the emergence of text-selectivity, while connectivity with frontal regions remains stable

We previously found that the intervention drove the emergence of the VWFA ^33^. To examine this effect spatially, we plotted the group probability maps of VWFA-1 and VWFA-2 within each timepoint. Each map shows the overlap of the individual participant’s ROI projected onto the average surface, such that each vertex in the map is color coded for the probability of being a part of the ROI in question. These maps (Figure 3A) visualize the emergence of the VWFA over time. In contrast, when looking at the connectivity map of the IFC, we observe no systematic change in its connectivity with the ventral surface. Notably, while these connectivity maps are not identical timepoint to timepoint, they never extend posteriorly towards VWFA-1, in line with the specific coupling between IFC and VWFA-2.

**Figure 3.**
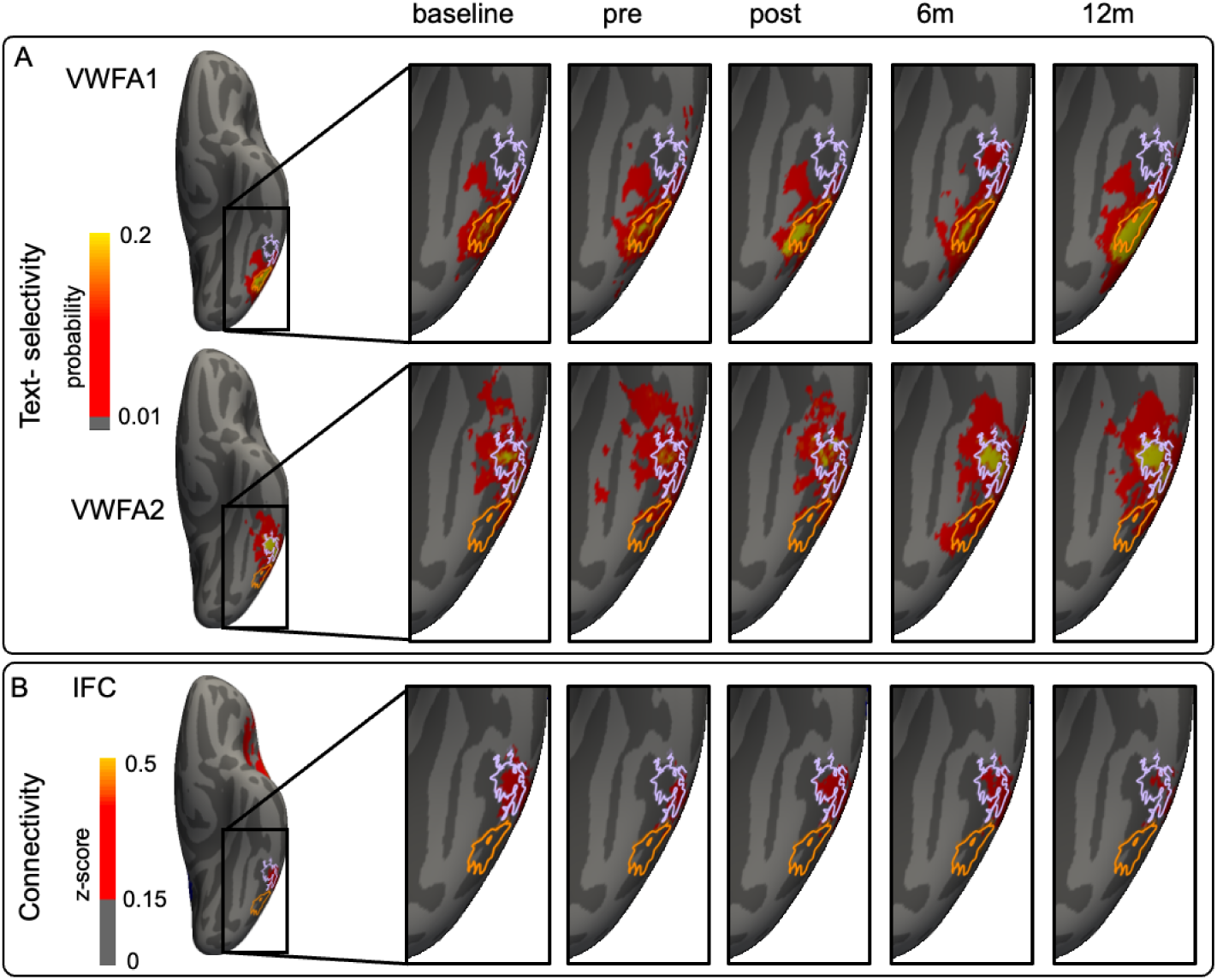
Functional connectivity remains stable while text selectivity increases. (A) Group probability maps of VWFA-1 (top) and VWFA-2 (bottom). Values in the map denote the probability of any vertex to include an individual ROI label (a value of 0.2 indicates that 20% of participants with usable data in that session had the ROI label in this location). Shown on the left are the probability maps of the combined ROIs (calculated over all available data for each child across timepoints), while insets show individual sessions. Contours denote the group VWFA-1 (orange) and VWFA-2 (lilac). (B) Average connectivity of the inferior frontal cortex (IFC) projected to average space and averaged across intervention participants. Shown on the left is the average connectivity map averaged across all sessions, while insets show individual sessions.

### The distance between VWFA-2 and VWFA-1 connectivity profiles is stable regardless of reading ability, text-selectivity and time

To go beyond connectivity with the IFC alone, we examined the connectivity patterns of VWFA-1 and VWFA-2 with the rest of the brain. To this end, we calculated a connectivity matrix using the Schaefer 200 atlas parcellation (see *Methods*), the IFC, and the four ventral ROIs (VWFA-1, VWFA-2, FFA-1, FFA-2). Using this matrix, we represented each ROI as a vector of connectivity values with the rest of the brain. We then calculated the cosine distance between VWFA-1 and VWFA-2 connectivity vectors, for each child and each timepoint in our data. Focusing on the pre-intervention timepoint, we found that the distance between VWFA-1 and VWFA-2 did not differ between typical readers and dyslexic readers (Figure 4B), or between children with or without a text-selective VWFA-2 (Figure 4C). We also observed that VWFA-2 connectivity profile was more similar to VWFA-1 than it was to FFA-2 and FFA-1 (Figure 4D). We then focused on the intervention group to ask whether following the intervention the VWFAs become more or less similar to each other in terms of functional connectivity with the rest of the brain.

**Figure 4.**
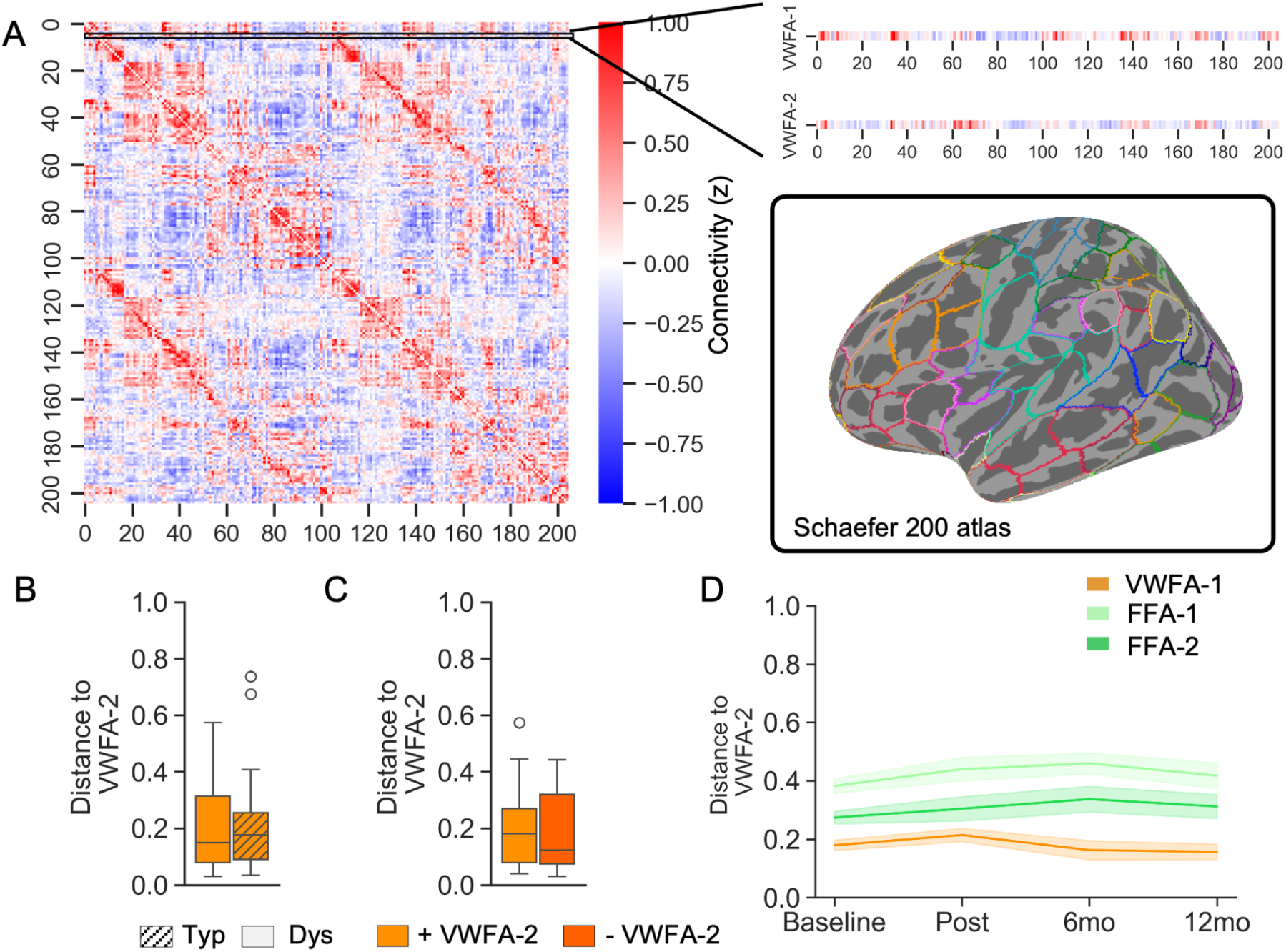
The distance in connectivity space between VWFA-2 and VWFA-1. (A) Whole brain connectivity matrix showing the connectivity strength (Fisher’s z transformed correlation values) between the ventral category-selective ROIs and the 200 Schaefer atlas parcellations (inset). Distance metrics are calculated between the correlation vectors representing the connectivity profiles of VWFA-1 and VWFA-2. (B) The distance between VWFA-2 and VWFA-1 is similar between children with dyslexia and typical readers (independent samples t-test: t = −0.64, p = 0.53). (C) Among children with dyslexia, the distance between VWFA-2 and VWFA-1 is similar between children with or without a text-selective VWFA-2 (t = 0.36, p = 0.72). (D) The distance between VWFA-2 and the other ventral ROIs remains stable over time in the intervention group. Lines denote the median and shaded areas denote one standard error of the mean.

We found that the distance between the connectivity profiles of VWFA-1 and VWFA-2 did not change over time in the intervention group (Figure 4D; time linear: *β* = −0.0003, p = 0.4565; time compared to baseline: *β* = −0.009, *β* = 0.0106, *β* = −0.0254 for post-intervention, 6 month and 12 month followup, respectively, all ps > 0.1). Similar results were obtained when we repeated the analyses using Pearson’s distance (1- Pearson’s correlation coefficient between the two vectors, see Supplementary Table S6).

### Connectivity predicts text-selectivity a year later

To investigate the notion that functional connectivity governs the location of VWFA-2, we examined the spatial relationship between functionally-defined ventral ROIs, and the connectivity pattern of the IFC. As a first step, we looked at each child’s seed-based connectivity map of the individually defined IFC. For this analysis, connectivity maps, as well as ROIs, were computed based on all available data for each child (collapsed across timepoints), to maximize signal to noise ratio (SNR). We calculated the Dice similarity coefficient between IFC connectivity map on the ventral surface and the ventral ROIs.

This analysis revealed that VWFA-2 showed greater spatial overlap with IFC connectivity compared with all other ROIs. This spatial overlap can be observed at the individual level (Figure 5A) as well as across the group (Figure 5B). We further confirmed the robustness of this relationship by repeating this analysis at different connectivity thresholds.

**Figure 5.**
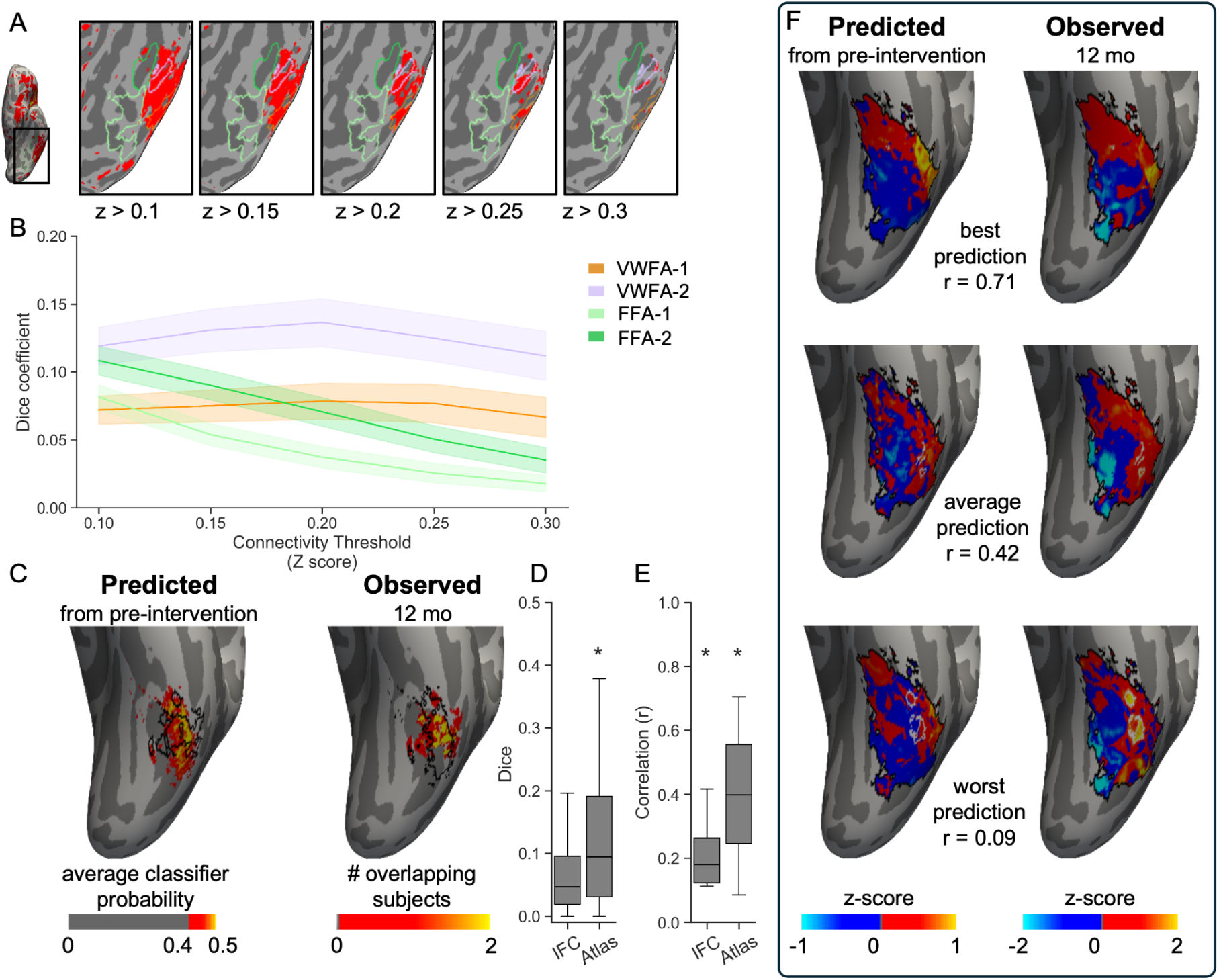
Functional connectivity predicts the location of VWFA-2. (A) Connectivity of the IFC overlaps VWFA-2 at the individual level. Shown are the manually labeled category selective regions for an individual subject, overlaid on the functional connectivity map of the IFC seed. Left to right, the images show the same map with gradually increasing connectivity thresholds. (B) Across different thresholds, VWFA-2 shows greater spatial overlap with IFC connectivity, compared with the other ventral ROIs. Each line represents the group average Dice similarity Coefficient (DC) across all subjects for whom individual ROIs were identified (N=65). Shaded areas denote one standard error of the mean. Across all thresholds, DC is higher between the IFC connectivity map and VWFA-2 compared with VWFA-1 (paired samples t-tests across the five thresholds: p = 0.003, p = 0.002, p = 0.004, p = 0.016, p = 0.039). (C) Functional connectivity data pre-intervention predicts VWFA-2 emergence a year later. Left- average group probability of VWFA-2, predicted using a classifier trained on whole brain functional connectivity data (the Atlas model). Black contour denotes the boundaries of the actual VWFA-2 observed in the held-out subjects. Right- Group map of the observed VWFA-2 in the held-out subjects. Black contour denotes the boundaries of the predicted VWFA-2 from the classifier. Thus, the predicted map is overlaid with the observed map contour and vice versa. Model significance was evaluated using a non-parametric permutation test (p < 1e-5). (D) DC between the measured and predicted VWFA-2 across individual subjects. Mean DC was significantly greater than zero for the Atlas model predictions (t_(9)_ = 3.118, p = 0.0123) and to a lesser degree for the IFC model predictions (t_(9)_ = 2.250, p = 0.0510). (E) Pearsons’ correlation across individuals between measured and predicted text-selectivity based on linear regression models. Mean r was significantly greater than zero for both the IFC model predictions (t_(9)_ = 2.491, p = 0.0343) and the Atlas model predictions (t_(9)_ = 6.222, p = 0.0002). (F) Comparison of observed and predicted text- selectivity in three individual subjects. Black contour denotes the boundaries of the search space for the models, defined as all vertices where any subject had a VWFA-2 at any given timepoint. White contours show the manually labeled VWFA-2 for each participant.

After observing this spatial coupling between connectivity and VWFA-2, we sought to test the hypothesis that functional connectivity before the intervention can predict where VWFA-2 will emerge after learning. To this end, we take advantage of a unique population within our sample: a subgroup of children with dyslexia who did not have a VWFA-2 prior to the intervention, but did develop one a year later ^33^. This included N=13 children, of whom 10 had sufficient functional connectivity data prior to the intervention (see *Methods* for data inclusion criteria). We designated this group as our held out test set, and trained models on all the available data from the other participants. Thus the training data included all available data concatenated across timepoints and participants, except for the held out participants. We first trained a classifier model on the IFC connectivity data, predicting in each vertex whether it was labeled VWFA-2 or not (the *IFC classifier* model). We then tested the model performance by applying the model to the connectivity data from the pre-intervention timepoint, to predict VWFA-2 in the final timepoint of the study, that is, the 12 month follow up. This model reached a f1 score of 0.109 (p = 0.0366, significance evaluated using a non-parametric permutation test permuting all the participants with 100000 iterations) and average precision score of 0.051 (p = 0.069). To test how well the model can predict VWFA-2 location within individuals, we also calculated the Dice similarity coefficient between the predicted and observed VWFA-2 in each child. The distribution of Dice values using this model was marginally greater than zero (t(9) = 2.250, p = 0.0510). We next fit a second model trained on richer data, this time using, in addition to the IFC connectivity vector, the connectivity of the 200 parcellations from the Schaefer Atlas (the *Atlas classifier* model). When evaluating the Atlas classifier model on the held out data, it reached f1 score of 0.153 (p = 0.0009, permutation test) and average precision of 0.078 (p = 0.0089, permutation test). Compared with the IFC classifier, the Atlas classifier performed much better in predicting VWFA-2 location in individual children (t_(9)_ = 3.118, p = 0.0123; Figure 5).

Lastly, we wanted to test whether the same data can predict the distributed response to text, rather than a binary classification of VWFA-2. This allows us to increase the size of the training data, since the data of children who did not have a VWFA-2 could not contribute to the classifier model, but could be used in the regression model predicting distributed text-selectivity. In addition, this analysis is less dependent on the specific criteria used to define the VWFA-2. Using the same predictors as before, we now trained an Elastic Net linear regression model to predict each vertex’s selectivity to text. The IFC regression model achieved a correlation value of r = 0.1596, p = 0.0675, and R^2^ score of 0.0214, p < e10-5 (permutation test). Including the entire connectivity matrix improved model performance to r = 0.413 and R^2^ score 0.1704, both p < e10-5 (permutation test). Across individual subjects, the correlation between the predicted and observed text-selectivity was significantly greater than zero for both the IFC regression model (t_(9)_ = 2.491, p = 0.0343) and the Atlas regression model (t_(9)_ = 6.222, p = 0.0002).

Together these results show that the model learned a relationship between functional connectivity and text-selectivity, which generalized to unseen subjects. Remarkably, the model was able to predict text-selectivity a year later from functional connectivity prior to the intervention.

## Discussion

Here we combined an educational intervention with longitudinal neuroimaging in a unique population of children with dyslexia, to ask whether there is a connectivity blueprint that predates the ability to read proficiently, and scaffolds the development of visual regions dedicated to processing words. Our key finding is that functional connectivity precedes and predicts the emergence of text-selective cortex. We show this in two complementary ways, first predicting the location of VWFA-2 in those who lacked the region prior to intervention, then predicting the distributed text-selective responses in VOTC. Remarkably, models were able to generalize and predict the emergence of text-selective responses in an unseen group of children who did not have a VWFA-2 at the beginning of the study. Our longitudinal data provide compelling support for the connectivity hypothesis ^3,10^ by showing that connectivity precedes functional specialization, and that this relationship can be reliably measured in individual children. While studies in adults have shown that functional connectivity can predict category selective regions in VOTC ^16,19^, this correspondence was only ever shown between tasks collected at the same timepoint, leaving open the developmental question of how this correspondence evolves, and whether there is a temporal precedence. A couple of longitudinal studies have shown that structural connectivity, as measured with diffusion MRI, can predict the location of category selective regions in children years later ^40,41^. To the best of our knowledge, our study is the first to show that functional connectivity serves as a blueprint that predates and predicts the emergence of category selectivity over the scale of months and over the course of development.

One view of functional connectivity holds that the functional coupling between distant brain regions forms via Hebbian-like learning principles, that is, stems from the joint activation of distant regions. The idea is that when distant brain regions are frequently co-activated during a task, they become synchronized at rest as well. Under this assumption, the strength of functional connectivity between regions reflects their history of co-activation. This led to suggestions that resting-state functional connectivity can be a promising indicator of learning and plasticity ^42,43^. This view has been supported by several studies that found changes in functional connectivity strength between specific regions following learning in adults ^44–46^. In the current study we did not observe evidence for such learning-induced plasticity in functional connectivity of the VWFA. Instead, our findings are more in line with the notion of functional connectivity as a stable trait or a blueprint for functional specialization ^47^.

Recent precision imaging studies collecting large amounts of data within individuals are leading to a view of functional networks as individual fingerprints that remain stable over time ^48,49^ and vary between individuals ^50^. Our current data corroborate this view of connectivity as an individualized innate functional architecture that allows brain regions to differentiate into specialized, functional modules.

Our findings contribute to the broader discussion on the role of experience and learning in shaping brain development. Research has sought to understand which brain systems are plastic, and what are the constraints on such plasticity in typical and clinical populations ^51^. In other words, what characteristics of the brain are innate and can serve as bio markers, and which are malleable to change with experience and can be an indicator of remediation? Learning to read provides a unique window into these questions, as a skill that involves a well-described network, but requires extensive training to develop. The VWFA was shown to have privileged connectivity with language regions in prereading (preverbal) infants ^20^, as well as in congenitally blind adults ^52^. It was thus suggested that functional connectivity is innate and does not require any sensory experience to develop. On the other hand, other studies have noted that in the congenitally blind, functional connectivity patterns are more diverse and show extensive reorganization, suggesting that visual experience actually is required to impose consistency on the preexisting network ^23,53,54^. These discrepancies have left many open questions about the effects of experience and learning on brain plasticity and reorganization. Our current investigation provides critical evidence for this debate by providing longitudinal data that go beyond a single snapshot of functional connectivity. Rather, our longitudinal design, combined with the intervention which systematically manipulated the learning environment, bring us a step closer to understanding the causal relationships between learning experience, functional specialization, and connectivity.

In sum, we provide here developmental evidence for the evolving relationship between functional connectivity, and text-selectivity. The intervention design allows us to portray the emerging VWFA-2 in children with dyslexia who undergo an intensive, controlled, learning experience. We show that while cortical responses to text increase with intervention, functional connectivity of the same regions is already in place before the intervention, and remains stable. Further, functional connectivity can predict text-selectivity a year later, providing evidence for the temporal precedence of functional connectivity. Our findings support a view of functional connectivity as an innate organizational principle of the cortex that scaffolds the subsequent emergence of functional regions following learning. Future research is needed for a precise characterization of how different learning experiences modulate this process, and what are the critical ingredients required to drive functional change and emergent behaviors like literacy. Together, our findings provide a nuanced understanding of how learning shapes the brain’s reading circuitry, and provide new insights into the interplay between connectivity, function, experience and behavior.

## Methods

### Participants

Forty-four children with dyslexia participated in the *Seeing Stars* 8-week reading intervention program delivered by Lindamood-Bell ^55^. Children were scanned up to 5 timepoints over the course of a year: 4-8 weeks before the intervention, immediately before the intervention, immediately after the intervention, 6 months later and 12 months later. The full timeline of the study, screening procedures and demographic characteristics of the sample have been reported in ^33^, and the key details are repeated here for convenience. In addition to the intervention participants, 43 children were scanned at similar timepoints without participating in the intervention. Of these, 24 children were typical readers and 19 children were struggling readers who enrolled to receive the intervention after completing the study. Children in the no-intervention group started the study with the pre-intervention visit, that is, completed 4 timepoints over the course of a year. The study was approved by the Stanford School of Medicine Institutional Review Board, and participants were compensated for their time. All participants provided assent and their parents provided written consent prior to participation.

### MRI data acquisition

In each visit, participants completed a functional category localizer that was used to identify their text-selective responses, and a movie watching scan that was used for functional connectivity analyses. Diffusion and quantitative scans were also collected as part of a large longitudinal study and are not part of the current analysis. In each visit the scanning session was typically split to two sessions with a break in between to allow participants to rest. Usually the localizer was collected in the first part and the movie scan was collected in the second part of the visit, but this order was flexible depending on the participant’s engagement.

Participants were scanned using a General Electric Sigma MR750 3T scanner at Stanford University’s Center for Cognitive and Neurobiological Imaging (CNI). During their first visit participants practiced the task and got acclimated to the scanner in a child-friendly mock MRI.

#### Category localizer

Functional runs were collected using a gradient echo EPI sequence with a multiband factor of 3, ensuring whole-brain coverage across 51 slices. The acquisition parameters included a TR of 1.19s, a TE of 30 ms, and a flip angle of 62, resulting in a spatial resolution of 2.4 mm³ isotropic voxels. Each run consisted of 232 frames and lasted 4 minutes and 36 seconds. In each visit participants completed four localizer runs, alternating between one-back and fixation color detection on the same visual stimuli. Data exclusion and analysis of the localizer data have been reported extensively in ^33,39^.

#### Passive movie watching

Two functional runs were collected using a gradient echo EPI sequence while the participants were watching a nature movie with no audio or text. The acquisition parameters included a TR of 820ms, a TE of 30 ms, and a flip angle of 54 with a hyperband factor of 6, resulting in a spatial resolution of 2.4 mm³ isotropic voxels and 66 slices. Each run included 375 frames, totalling 5 minutes and 8 seconds. Fieldmaps with opposing directions (pe0 and pe1) were collected with the same prescription prior to the functional runs to allow for EPI distortion correction (separate fieldmaps were collected for the category localizer and for the movie runs due to the different coverage). Lastly, a high-resolution T1-weighted anatomical scan was acquired with a spatial resolution of 0.9 mm³ isotropic voxels.

### MRI data preprocessing

Data were preprocessed using *fmriprep* version 23.1.3 ^56^. Confounds extracted from fmriprep were used as regressors for signal cleaning (see below). All analyses were carried out using Nilearn v0.10.1 and visualized with Freeview v7.3.2 (https://surfer.nmr.mgh.harvard.edu/).

### Data inclusion

Resting state runs were flagged for exclusion if they had a mean framewise displacement (FD) greater than 0.5 mm, or if more than 20% of volumes had a FD greater than 0.5 mm. We only included sessions with two complete usable runs. Additional 10 sessions were excluded due to a technical artifact in the scanning facility. This resulted in a total of 301 sessions in 87 children.

### Region of interest definition

In each visit participants completed 4 runs of a visual functional localizer as described in ^33,34,39^. In brief, participants viewed text and non-text (faces, limbs, objects) visual stimuli and performed a one-back task or a fixation color detection task alternately. To identify text-selective responses in individual participants, we used the contrast Text> all other stimuli, thresholded at t > 3. Text-selectivity maps were generated based on all data available for each subject (collapsed across timepoints) to maximize SNR and ensure the same ROIs are being used across timepoints. This is important to avoid conflating functional connectivity with the observed changes in VWFA size over the intervention ^33^. Then, VWFA-1 and VWFA-2 were hand drawn using anatomical landmarks as described in ^33^. In addition, the same anatomical guidelines were used to define the ROIs separately at each timepoint. The same procedure was used to identify face-selective ROIs, namely FFA-1 and FFA-2, as control regions. Lastly, we identified text-selective responses in the frontal lobe as described in ^34^. For the current analyses, we created a lenient frontal lobe text-selective ROI that included the union of text-selective activations in the inferior frontal gyrus and precentral gyrus. These native ROIs allowed us to (1) investigate functional connectivity specifically between distant text-selective regions in each individual, and (2) identify a unique population of children who did not show text-selectivity in the cortical area of VWFA-2.

To explore functional connectivity in participants who didn’t show text selectivity (the *-VWFA2* group), we created a second set of ROIs that allowed us to carry out analyses in template space. To this end, we projected all the ROIs from native space to the template *fsaverage* space and created a group probability map where each vertex represented the probability of any subject having the ROI in that location. We thresholded these probability maps at p > 0.15 to create group template ROIs (following the same procedure as in ^32^). Since the FFAs resulting from this procedure were much larger than the VWFAs (due to the known higher variability in VWFA location), we enforced a uniform size on the ventral ROIs to avoid size differences biasing the results. To this end, the ventral ROIs were limited to equal sizes by taking the number of vertices of the smallest ROI size (n), selecting the top n vertices from all other ROIs, and removing overlapping vertices.

### Functional connectivity data processing

Preprocessed data were cleaned using Nilearn *signal.clean* function [Nilearn 0.10.1; ^57^]. The first 6 frames of each run were discarded to allow the signal to achieve steady state. Confound regression was performed with six motion regressors, white matter signal, cerebro-spinal fluid signal, global signal, and their first derivatives, resulting in 18 regressors. Volumes where motion exceeded framewise displacement (FD) of 0.5mm were flagged for scrubbing and included as additional regressors. In addition, we applied a bandpass butterworth filter in the range [0.008-0.1Hz], detrending and standardization of the signal, as implemented in *signal.clean.* To create connectivity maps, we calculated the mean time series in a given ROI and calculated Pearson’s correlation between that seed and the signal timeseries of each vertex on the cortical surface. Correlation coefficients were transformed to z-values using the Fisher transform. The same pipeline was applied to data in native space and in fsaverage space, which were used for different analyses as reported in the result section.

### Statistical analysis

To assess differences in connectivity between VWFA-1 and VWFA-2, we subtracted the VWFA-2 connectivity map from that of VWFA-1 (such that negative values indicate higher connectivity with VWFA-2). To directly test the hypothesis that VWFA-2 is more strongly connected to specific language regions in the frontal lobe, we ran an ROI- to-ROI analysis where we quantified the connectivity strength between pairs of ROIs. Specifically, we calculate Pearson’s correlation coefficient between the timeseries of VWFA-1 and IFC-text, and VWFA-2 and IFC-text. These connectivity values were then entered into linear mixed effect (LME) models using the *lme4* package in R ^58^. We first tested whether connectivity strength between IFC and VWFA-2 was greater compared with other ventral ROIs, with an interaction term of reading group, accounting for participant’s age and in-scanner motion:

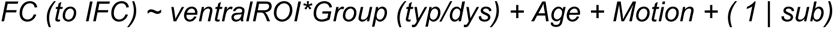

We next ran a similar model, this time testing whether connectivity strength was different between participants who had a text-selective VWFA-2 and those who didn’t show text-selectivity in this area:

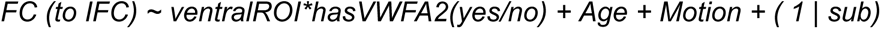

Lastly, to investigate whether functional connectivity changed with time, we ran a series of longitudinal models testing whether connectivity strength between IFC and VWFA-2 or VWFA-1 was modulated by time (number of weeks since pre-intervention visit). To capture potential non-linear time effects, we repeated this analysis while treating timepoint as a categorical variable, such that each visit was compared to the baseline, coded as the average of the two pre-intervention visits. Unless otherwise noted, VWFA-2 was set as the reference level in all models.

### Distance analysis

In addition to the pre-defined ROIs, we investigate whole brain connectivity using the 200 cortical parcellations of the Schaefer Atlas ^59^. We created a connectivity matrix of the 200 parcellations in addition to VWFA-1, VWFA-2, FFA-1, FFA-2 and IFC-text (matrix size 205 * 205). Each cell in the matrix represents the Fisher transformed Pearson’s correlation between ROI pairs. This matrix was used to evaluate the distance in connectivity space between VWFA-1 and VWFA-2. The distance was defined as the Cosine distance between the connectivity vector of VWFA-1, and the connectivity vector of VWFA-2 (see Figure 4). Two tailed independent samples t-tests were used to test for group differences pre-intervention, and LME models were used to evaluate longitudinal changes in the distance between VWFA-1 and VWFA-2.

### Spatial overlap

We used the Dice similarity coefficient (DC) to quantify the spatial overlap between the connectivity of IFC on the ventral surface and each of the four ventral ROIs. For this analysis, we used connectivity maps and ROIs created based on all available data for each subject, averaged across timepoints, to maximize SNR. We constrained the calculation to an anatomical VOTC region and used a threshold to capture vertices showing high IFC connectivity. Then, in each subject, we calculated the DC as the number of overlapping vertices, divided by the sum of the number of vertices in the thresholded FC map and the number of vertices in the ROI in question. As a sensitivity analysis, we repeated this calculation for different thresholds of the IFC connectivity map.

### Prediction modeling

To test the hypothesis that functional connectivity governs the emergence of functional regions, we took a predictive modeling approach using functional connectivity data to predict text-selectivity. We fit two types of models: a classifier to determine whether each vertex belongs to VWFA-2 or not, and a regression model to predict each vertex’s degree of text-selectivity. For each model, our held out dataset consisted of a unique group of subjects that did not have a VWFA-2 at their pre-intervention session but did develop it at the last timepoint. We refer to these subjects (N = 13) as the *emerging VWFA-2* group. We trained each model on all available data from the rest of the participants, collapsing across all available timepoints. Then we evaluated the model performance on the held out participants, by running it on their connectivity data from the pre-intervention session, predicting their VWFA-2 in the 12 months follow up.

The predictor data for all models was functional connectivity: we first used the connectivity map of the IFC-text as the predictor variable, given the observed correspondence between IFC connectivity and VWFA-2 (see Figure 5). To explore whether whole-brain connectivity further improves the ability to predict text responses, we used the Schaefer atlas parcellations as seeds and created a predictor matrix of 201 regions (200 parcellations + IFC-text) by the number of vertices in the left hemisphere. Thus, in this matrix, each vertex is represented by a vector of connectivity values with 201 regions. We constrained our analyses to a search space defined as all vertices where any subject had a VWFA-2 identified at any given timepoint (3512 vertices). All data were z-scored to control for individual differences in mean connectivity values before being entered into the model. All models were trained and tested using scikit-learn version 1.6.1 ^60^.

We first fit a gradient-boosted random forest model (XGBoost, version 2.1.4; ^61^) to predict VWFA-2 (a binary vector). Before running each model, we performed hyperparameter tuning using Bayesian optimization on the training data, with 3 cross-validation folds and 50 iterations, using the *skopt* package, version 0.10.2 ^62^. We evaluated model performance as the f1 score, defined as the harmonic mean of precision and recall, and the average precision score. We evaluated significance by running a non-parametric permutation test with 100000 iterations, where we randomly permuted the subjects in the outcome vector and recalculated the performance metric (f1 and average precision) between the predicted outcome and the permuted outcome.

Next, we fit an Elastic Net linear regression to predict text-selectivity as a continuous variable. This approach has the advantage that it provides more nuanced prediction compared with the binary classification, is unaffected by arbitrary threshold selection used for ROI definition, and includes more data as it allows to predict low values of text-selectivity in participants without a VWFA-2. We evaluated model predictions using the coefficient of determination (R2) and Pearson’s correlation between the predicted and measured responses. Significance was evaluated in the same manner using a non-parametric permutation test, comparing model performance to a null distribution of 100000 random permutations.

## Acknowledgements

This work was supported by NICHD R01-HD095861 to JDY. We thank the children and families who participated in the study.

## Notes

### Competing Interest Statement

The authors have declared no competing interest.

## References

1. Op de Beeck, H. P., Pillet, I. & Ritchie, J. B. Factors Determining Where Category-Selective Areas Emerge in Visual Cortex. Trends Cogn. Sci. 23, 784–797 (2019).

2. Grill-Spector, K. & Weiner, K. S. The functional architecture of the ventral temporal cortex and its role in categorization. Nat. Rev. Neurosci. 15, 536–548 (2014).

3. Hannagan, T., Amedi, A., Cohen, L., Dehaene-Lambertz, G. & Dehaene, S. Origins of the specialization for letters and numbers in ventral occipitotemporal cortex. Trends Cogn. Sci. 19, 374–382 (2015).

4. Hasson, U., Levy, I., Behrmann, M., Hendler, T. & Malach, R. Eccentricity bias as an organizing principle for human high-order object areas. Neuron 34, 479–490 (2002).

5. Levy, I., Hasson, U., Avidan, G., Hendler, T. & Malach, R. Center-periphery organization of human object areas. Nat. Neurosci. 4, 533–539 (2001).

6. Ellis, C. T. et al. Retinotopic organization of visual cortex in human infants. Neuron 109, 2616–2626.e6 (2021).

7. Long, B., Yu, C.-P. & Konkle, T. Mid-level visual features underlie the high-level categorical organization of the ventral stream. Proc. Natl. Acad. Sci. U. S. A. 115, E9015–E9024 (2018).

8. Bao, P., She, L., McGill, M. & Tsao, D. Y. A map of object space in primate inferotemporal cortex. Nature 583, 103–108 (2020).

9. Mahon, B. Z. & Caramazza, A. What drives the organization of object knowledge in the brain? Trends Cogn. Sci. 15, 97–103 (2011).

10. Passingham, R. E., Stephan, K. E. & Kötter, R. The anatomical basis of functional localization in the cortex. Nat. Rev. Neurosci. 3, 606–616 (2002).

11. Smith, S. M. et al. Correspondence of the brain’s functional architecture during activation and rest. Proc. Natl. Acad. Sci. U. S. A. 106, 13040–13045 (2009).

12. Bernstein-Eliav, M. & Tavor, I. The prediction of brain activity from connectivity: Advances and applications. Neuroscientist 30, 367–377 (2024).

13. Tavor, I. et al. Task-free MRI predicts individual differences in brain activity during task performance. Science 352, 216–220 (2016).

14. Hiersche, K. J., Saygin, Z. M. & Osher, D. E. Connectivity and function are coupled across cognitive domains throughout the brain. Netw. Neurosci. 10, 80–92 (2026).

15. Cole, M. W., Ito, T., Bassett, D. S. & Schultz, D. H. Activity flow over resting-state networks shapes cognitive task activations. Nat. Neurosci. 19, 1718–1726 (2016).

16. Molloy, M. F., Saygin, Z. M. & Osher, D. E. Predicting high-level visual areas in the absence of task fMRI. Sci. Rep. 14, 11376 (2024).

17. Conrad, B. N., Pollack, C., Yeo, D. J. & Price, G. R. Structural and functional connectivity of the inferior temporal numeral area. Cereb. Cortex 33, 6152–6170 (2023).

18. Hutchison, R. M., Culham, J. C., Everling, S., Flanagan, J. R. & Gallivan, J. P. Distinct and distributed functional connectivity patterns across cortex reflect the domain-specific constraints of object, face, scene, body, and tool category-selective modules in the ventral visual pathway. Neuroimage 96, 216–236 (2014).

19. Chen, Q., Garcea, F. E., Almeida, J. & Mahon, B. Z. Connectivity-based constraints on category-specificity in the ventral object processing pathway. Neuropsychologia 105, 184–196 (2017).

20. Li, J., Osher, D. E., Hansen, H. A. & Saygin, Z. M. Innate connectivity patterns drive the development of the visual word form area. Sci. Rep. 10, 18039 (2020).

21. Kamps, F. S., Hendrix, C. L., Brennan, P. A. & Dilks, D. D. Connectivity at the origins of domain specificity in the cortical face and place networks. Proceedings of the National Academy of Sciences 117, 6163–6169 (2020).

22. Lesinger, K. et al. Functional connectivity of the human face network exhibits right hemispheric lateralization from infancy to adulthood. Sci. Rep. 13, 20831 (2023).

23. Striem-Amit, E. et al. Functional connectivity of visual cortex in the blind follows retinotopic organization principles. Brain 138, 1679–1695 (2015).

24. Striem-Amit, E. et al. Topographical functional connectivity patterns exist in the congenitally, prelingually deaf. Sci. Rep. 6, 29375 (2016).

25. Dehaene, S. & Cohen, L. Cultural recycling of cortical maps. Neuron 56, 384–398 (2007).

26. Cohen, L. et al. Language-specific tuning of visual cortex? Functional properties of the Visual Word Form Area. Brain 125, 1054–1069 (2002).

27. Dehaene, S. & Cohen, L. The unique role of the visual word form area in reading. Trends Cogn. Sci. 15, 254–262 (2011).

28. Lerma-Usabiaga, G., Carreiras, M. & Paz-Alonso, P. M. Converging evidence for functional and structural segregation within the left ventral occipitotemporal cortex in reading. Proceedings of the National Academy of Sciences 115, E9981–E9990 (2018).

29. White, A. L., Palmer, J., Boynton, G. M. & Yeatman, J. D. Parallel spatial channels converge at a bottleneck in anterior word-selective cortex. Proceedings of the National Academy of Sciences 116, 10087–10096 (2019).

30. Yeatman, J. D. & White, A. L. Reading: The Confluence of Vision and Language. Annual Review of Vision Science 7, 487–517 (2021).

31. Caffarra, S., Karipidis, I. I., Yablonski, M. & Yeatman, J. D. Anatomy and physiology of word-selective visual cortex: from visual features to lexical processing. Brain Struct. Funct. 226, 3051–3065 (2021).

32. Yablonski, M., Karipidis, I. I., Kubota, E. & Yeatman, J. D. The transition from vision to language: Distinct patterns of functional connectivity for subregions of the visual word form area. Hum. Brain Mapp. 45, e26655 (2024).

33. Mitchell, J. L. et al. The balance between stability and plasticity of the visual word form area in dyslexia. Nat. Commun. (2025) doi:10.1038/s41467-025-67054-3.

34. Stone, H. L. et al. Anatomically distinct regions in the inferior frontal cortex are modulated by task and reading skill. J. Neurosci. 45, (2025).

35. Finn, E. S. & Bandettini, P. A. Movie-watching outperforms rest for functional connectivity-based prediction of behavior. Neuroimage 235, 117963 (2021).

36. Gal, S., Coldham, Y., Tik, N., Bernstein-Eliav, M. & Tavor, I. Act natural: Functional connectivity from naturalistic stimuli fMRI outperforms resting-state in predicting brain activity. Neuroimage 258, 119359 (2022).

37. Vanderwal, T., Eilbott, J. & Castellanos, F. X. Movies in the magnet: Naturalistic paradigms in developmental functional neuroimaging. Dev. Cogn. Neurosci. 36, 100600 (2019).

38. Glezer, L. S. & Riesenhuber, M. Individual Variability in Location Impacts Orthographic Selectivity in the ‘Visual Word Form Area’. Journal of Neuroscience 33, 11221–11226 (2013).

39. Mitchell, J. L., Jimenez, M., Stone, H. L., Yablonski, M. & Yeatman, J. D. Visual Word Form Area demonstrates individual and task-agnostic consistency but inter-individual variability. Dev. Cogn. Neurosci. 79, 101703 (2026).

40. Saygin, Z. M. et al. Anatomical connectivity patterns predict face selectivity in the fusiform gyrus. Nat. Neurosci. 15, 321–327 (2012).

41. Saygin, Z. M. et al. Connectivity precedes function in the development of the visual word form area. Nat. Neurosci. 19, 1250–1255 (2016).

42. Guerra-Carrillo, B., Mackey, A. P. & Bunge, S. A. Resting-state fMRI: a window into human brain plasticity. Neuroscientist 20, 522–533 (2014).

43. Kelly, C. & Castellanos, F. X. Strengthening connections: functional connectivity and brain plasticity. Neuropsychol. Rev. 24, 63–76 (2014).

44. Mackey, A. P., Miller Singley, A. T. & Bunge, S. A. Intensive reasoning training alters patterns of brain connectivity at rest. J. Neurosci. 33, 4796–4803 (2013).

45. Harmelech, T., Preminger, S., Wertman, E. & Malach, R. The day-after effect: long term, Hebbian-like restructuring of resting-state fMRI patterns induced by a single epoch of cortical activation. J. Neurosci. 33, 9488–9497 (2013).

46. Lewis, C. M., Baldassarre, A., Committeri, G., Romani, G. L. & Corbetta, M. Learning sculpts the spontaneous activity of the resting human brain. Proc. Natl. Acad. Sci. U. S. A. 106, 17558–17563 (2009).

47. Gratton, C. et al. Functional brain networks are dominated by stable group and individual factors, not cognitive or daily variation. Neuron 98, 439–452.e5 (2018).

48. Amaral, L., Thomas, P., Amedi, A. & Striem-Amit, E. Longitudinal stability of individual brain plasticity patterns in blindness. Proc. Natl. Acad. Sci. U. S. A. 121, e2320251121 (2024).

49. Hu, D. et al. Existence of functional connectome fingerprint during infancy and its stability over months. J. Neurosci. 42, 377–389 (2022).

50. Braga, R. M. & Buckner, R. L. Parallel interdigitated distributed networks within the individual estimated by intrinsic functional connectivity. Neuron 95, 457–471.e5 (2017).

51. Wandell, B. A. & Smirnakis, S. M. Plasticity and stability of visual field maps in adult primary visual cortex. Nat. Rev. Neurosci. 10, 873–884 (2009).

52. Abboud, S., Maidenbaum, S., Dehaene, S. & Amedi, A. A number-form area in the blind. Nat. Commun. 6, 6026 (2015).

53. Butt, O. H., Benson, N. C., Datta, R. & Aguirre, G. K. The fine-scale functional correlation of striate cortex in sighted and blind people. J. Neurosci. 33, 16209–16219 (2013).

54. Sen, S. et al. The role of visual experience in individual differences of brain connectivity. J. Neurosci. 42, 5070–5084 (2022).

55. Bell, N. Seeing Stars: Symbol Imagery for Phonological and Orthographic Processing in Reading and Spelling. (2013).

56. Esteban, O. et al. fMRIPrep: a robust preprocessing pipeline for functional MRI. Nat. Methods 16, 111–116 (2019).

57. Nilearn contributors et al. Nilearn. (Zenodo, 2024). doi:10.5281/ZENODO.10948303.

58. Bates, D., Mächler, M., Bolker, B. & Walker, S. Fitting Linear Mixed-Effects Models Using lme4. J. Stat. Softw. 67, 1–48 (2015).

59. Schaefer, A. et al. Local-global parcellation of the human cerebral cortex from intrinsic functional connectivity MRI. Cereb. Cortex 28, 3095–3114 (2018).

60. Pedregosa, F. et al. Scikit-learn: Machine learning in Python. The Journal of machine Learning research 12, 2825–2830 (2011).

61. Chen, T. & Guestrin, C. Xgboost: A scalable tree boosting system. Proceedings of the 22nd acm sigkdd international conference on knowledge discovery and data mining 785–794 (2016).

62. Louppe, G. & Kumar, M. Bayesian optimization with skopt — scikit-optimize 0.8.1 documentation. (2016).

