## Supplementary data for "Functional connectivity scaffolds the emergence of the visual word form area"

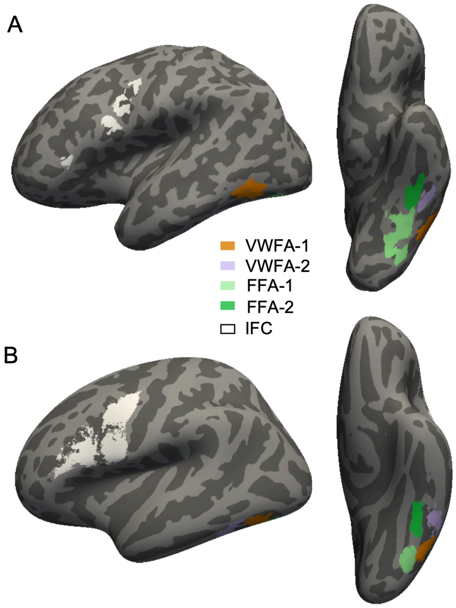


**Figure S1. Regions of interest used as seeds.** A) Individually defined ROIs based on text-selectivity in a functional localizer (response to Text> all other stimuli, thresholded at t> 3 for ventral regions and t>2 for frontal regions). Shown is the cortical surface of an individual participant (M, age 12y). B) Group ROIs based on probability across subjects, in fsaverage space. Individual ROIs were projected to fsaverage space, then the resulting probability map was thresholded at $p > 0.15$ (same procedure as in Yablonski et al., 2024). Ventral ROIs were limited to equal sizes by taking the number of vertices of the smallest ROI size (n) and selecting the top n vertices from all other ROIs.

| FC (to IFC) ~ ventral ROI + Age + Motion + ( 1 \| sub) | | | | | | | | | | |
|---|---|---|---|---|---|---|---|---|---|---|
| | Dyslexia readers (N= 53) | | | | | Typical readers (N = 19) | | | | |
| | β | se | t | df | p | β | se | t | df | p |
| Intercept (VWFA2) | 0.48 | 0.21 | 2.30 | 50.63 | **0.026** | 0.48 | 0.40 | 1.21 | 16.13 | 0.245 |
| VWFA1 | -0.14 | 0.03 | -5.34 | 156.00 | **0.000** | -0.20 | 0.04 | -4.91 | 54.00 | **0.000** |
| FFA1 | -0.32 | 0.03 | -12.09 | 156.00 | **0.000** | -0.38 | 0.04 | -9.22 | 54.00 | **0.000** |
| FFA2 | -0.21 | 0.03 | -7.83 | 156.00 | **0.000** | -0.30 | 0.04 | -7.29 | 54.00 | **0.000** |
| Age | -0.01 | 0.02 | -0.38 | 50.00 | 0.702 | -0.02 | 0.04 | -0.46 | 16.00 | 0.654 |
| Motion | -0.42 | 0.34 | -1.22 | 50.00 | 0.229 | 0.79 | 0.77 | 1.02 | 16.00 | 0.321 |

**Table S1.** Linear mixed effect models in the pre-intervention timepoint, comparing functional connectivity between each of the ventral ROIs (VWFA-1, VWFA-2, FFA-1, FFA-2) and the inferior frontal cortex (IFC), using template ROIs. Models were run separately in children with dyslexia (N=53) and typical readers (N=19). Age- number of years at the pre-intervention visit. Motion- mean framewise displacement (mm). VWFA-2 was set as the reference level.

| FC (to IFC) ~ ventral ROI + Age + Motion + ( 1 \| sub) | | | | | | | | | | |
|---|---|---|---|---|---|---|---|---|---|---|
| | With VWFA2 (N= 33) | | | | | Without VWFA2 (N = 20) | | | | |
| | β | se | t | df | p | β | se | t | df | p |
| Intercept (VWFA2) | 0.41 | 0.30 | 1.39 | 30 | 0.176 | 0.49 | 0.31 | 1.56 | 17 | 0.137 |
| VWFA1 | -0.14 | 0.03 | -4.44 | 96 | **0.000** | -0.14 | 0.05 | -2.97 | 57 | **0.004** |
| FFA1 | -0.32 | 0.03 | -10.00 | 96 | **0.000** | -0.32 | 0.05 | -6.76 | 57 | **0.000** |
| FFA2 | -0.22 | 0.03 | -6.65 | 96 | **0.000** | -0.20 | 0.05 | -4.19 | 57 | **0.000** |
| Age | 0.00 | 0.03 | 0.08 | 30 | 0.940 | -0.02 | 0.03 | -0.64 | 17 | 0.531 |
| Motion | -0.59 | 0.42 | -1.41 | 30 | 0.168 | 0.12 | 0.64 | 0.18 | 17 | 0.857 |

**Table S2.** Linear mixed effect models in the pre-intervention timepoint, comparing functional connectivity between each of the ventral ROIs (VWFA-1, VWFA-2, FFA-1, FFA-2) and the inferior frontal cortex (IFC), using template ROIs. Models were run separately in children where a VWFA-2 was identified based on text selectivity (N=33) and children where no VWFA-2 was identified (N=20). Age- number of years at the pre-intervention visit. Motion- mean framewise displacement (mm). VWFA-2 was set as the reference level.

| FC (to IFC) ~ ventral ROI + Age + Motion + ( 1 \| sub) | | | | | | | | | | | | | | | | | | | | |
|---|---|---|---|---|---|---|---|---|---|---|---|---|---|---|---|---|---|---|---|---|
| | pre-intervention | | | | | post- intervention | | | | | 6 months | | | | | 12 months | | | | |
| | β | se | t | df | p | β | se | t | df | p | β | se | t | df | p | β | se | t | df | p |
| **Intercept (VWFA2)** | 0.36 | 0.30 | 1.19 | 34 | 0.2409 | 0.42 | 0.34 | 1.25 | 25 | 0.2222 | 0.52 | 0.26 | 1.99 | 33 | 0.0553 | 0.36 | 0.26 | 1.39 | 36 | 0.1742 |
| **VWFA1** | -0.13 | 0.03 | -3.68 | 108 | **0.0004** | -0.19 | 0.04 | -5.32 | 81 | **0.0000** | -0.14 | 0.04 | -3.88 | 105 | **0.0002** | -0.14 | 0.03 | -3.99 | 114 | **0.0001** |
| **FFA1** | -0.31 | 0.03 | -9.02 | 108 | **0.0000** | -0.35 | 0.04 | -9.59 | 81 | **0.0000** | -0.33 | 0.04 | -8.91 | 105 | **0.0000** | -0.34 | 0.03 | -9.89 | 114 | **0.0000** |
| **FFA2** | -0.22 | 0.03 | -6.28 | 108 | **0.0000** | -0.26 | 0.04 | -7.18 | 81 | **0.0000** | -0.24 | 0.04 | -6.36 | 105 | **0.0000** | -0.21 | 0.03 | -6.14 | 114 | **0.0000** |
| **Age** | 0.00 | 0.03 | 0.05 | 34 | 0.9570 | 0.00 | 0.03 | -0.06 | 25 | 0.9526 | -0.02 | 0.02 | -0.67 | 33 | 0.5086 | -0.01 | 0.02 | -0.36 | 36 | 0.7186 |
| **Motion** | -0.34 | 0.50 | -0.68 | 34 | 0.5035 | 0.08 | 0.50 | 0.15 | 25 | 0.8792 | -0.09 | 0.51 | -0.18 | 33 | 0.8605 | 0.48 | 0.40 | 1.21 | 36 | 0.2343 |

**Table S3.** Linear mixed effect models within each timepoint, comparing functional connectivity between each of the ventral ROIs (VWFA-1, VWFA-2, FFA-1, FFA-2) and the inferior frontal cortex (IFC), **using template ROIs**, in the intervention group (N=44). Age is years at the pre-intervention visit. Motion- mean framewise displacement (mm).

| FC (to IFC) ~ ventral ROI + Age + Motion + ( 1 \| sub) | | | | | | | | | | | | | | | | | | | | |
|---|---|---|---|---|---|---|---|---|---|---|---|---|---|---|---|---|---|---|---|---|
| | pre-intervention | | | | | post- intervention | | | | | 6 months | | | | | 12 months | | | | |
| | β | se | t | df | p | β | se | t | df | p | β | se | t | df | p | β | se | t | df | p |
| **Intercept (VWFA2)** | 0.15 | 0.25 | 0.61 | 36 | 0.5476 | -0.31 | 0.38 | -0.80 | 25 | 0.4303 | -0.01 | 0.29 | -0.04 | 34 | 0.9722 | -0.09 | 0.35 | -0.27 | 37 | 0.7889 |
| **VWFA1** | -0.18 | 0.04 | -4.02 | 99 | **0.0001** | -0.24 | 0.05 | -4.82 | 72 | **0.0000** | -0.15 | 0.05 | -3.35 | 98 | **0.0012** | -0.12 | 0.05 | -2.52 | 105 | **0.0134** |
| **FFA1** | -0.38 | 0.04 | -8.77 | 99 | **0.0000** | -0.36 | 0.05 | -7.63 | 72 | **0.0000** | -0.38 | 0.04 | -8.52 | 97 | **0.0000** | -0.33 | 0.05 | -7.33 | 104 | **0.0000** |
| **FFA2** | -0.32 | 0.04 | -7.35 | 99 | **0.0000** | -0.37 | 0.05 | -7.69 | 72 | **0.0000** | -0.35 | 0.04 | -7.87 | 97 | **0.0000** | -0.25 | 0.05 | -5.64 | 104 | **0.0000** |
| **Age** | 0.03 | 0.02 | 1.07 | 35 | 0.2904 | 0.06 | 0.04 | 1.81 | 25 | 0.0822 | 0.04 | 0.03 | 1.32 | 33 | 0.1945 | 0.03 | 0.03 | 1.02 | 36 | 0.3142 |
| **Motion** | -0.28 | 0.43 | -0.65 | 35 | 0.5171 | 0.37 | 0.57 | 0.66 | 25 | 0.5159 | -0.04 | 0.58 | -0.07 | 33 | 0.9476 | 0.29 | 0.53 | 0.54 | 36 | 0.5898 |

**Table S4.** Linear mixed effect models within each timepoint, comparing functional connectivity between each of the ventral ROIs (VWFA-1, VWFA-2, FFA-1, FFA-2) and the inferior frontal cortex (IFC), using **individually defined ROIs**, in the intervention group (N=44). Age is years at the pre-intervention visit. Motion- mean framewise displacement (mm).

| | FC (VWFA1-IFC) ~ Time (weeks) + Age + Motion + ( 1 \| sub) | | | | | FC (VWFA2-IFC) ~ Time (weeks) + Age + Motion + ( 1 \| sub) | | | | |
|---|---|---|---|---|---|---|---|---|---|---|
| | β | se | t | df | p | β | se | t | df | p |
| **(Intercept)** | -0.43 | 0.36 | -1.19 | 36.83 | 0.2404 | 0.55 | 0.41 | 1.34 | 32.44 | 0.1884 |
| **Time (weeks)** | 0.00 | 0.00 | 1.10 | 123.15 | 0.2718 | 0.00 | 0.00 | -1.62 | 106.08 | 0.1075 |
| **Age** | 0.05 | 0.03 | 1.55 | 34.76 | 0.1301 | -0.01 | 0.04 | -0.30 | 30.92 | 0.7666 |
| **Motion** | 0.19 | 0.30 | 0.63 | 148.39 | 0.5319 | -0.43 | 0.34 | -1.28 | 129.48 | 0.2027 |

| | FC (VWFA1-IFC) ~ Time (session) + Age + Motion + ( 1 \| sub) | | | | | FC (VWFA2-IFC) ~ Time (session) + Age + Motion + ( 1 \| sub) | | | | |
|---|---|---|---|---|---|---|---|---|---|---|
| | β | se | t | df | p | β | se | t | df | p |
| **(Intercept)** | -0.43 | 0.36 | -1.22 | 37.13 | 0.2317 | 0.54 | 0.41 | 1.32 | 32.73 | 0.1967 |
| **session: post** | 0.03 | 0.04 | 0.61 | 120.67 | 0.5414 | 0.02 | 0.05 | 0.39 | 104.49 | 0.7001 |
| **session: 6mo** | 0.03 | 0.04 | 0.86 | 120.74 | 0.3909 | -0.02 | 0.04 | -0.57 | 104.69 | 0.5706 |
| **session: 12mo** | 0.03 | 0.04 | 0.83 | 119.92 | 0.4097 | -0.07 | 0.04 | -1.69 | 103.80 | 0.0945 |
| **Age** | 0.05 | 0.03 | 1.56 | 34.74 | 0.1276 | -0.01 | 0.04 | -0.29 | 31.00 | 0.7727 |
| **Motion** | 0.19 | 0.30 | 0.64 | 145.76 | 0.5247 | -0.41 | 0.34 | -1.20 | 127.55 | 0.2311 |

**Table S5.** Linear mixed effect models within each timepoint, comparing functional connectivity between each of the ventral ROIs (VWFA-1, VWFA-2, FFA-1, FFA-2) and the inferior frontal cortex (IFC), using **individually defined ROIs**, in the intervention group (N=44). This table parallels Table 3 from the main text. Age- number of years at the pre-intervention visit. Motion- mean framewise displacement (mm).

| | Distance (VWFA2-VWFA1) ~ Time (weeks) + Age + Motion + ( 1 \| sub | | | | | | | | | |
|---|---|---|---|---|---|---|---|---|---|---|
| | Cosine Distance | | | | | Pearson's distance | | | | |
| | β | se | t | df | p | β | se | t | df | p |
| **(Intercept)** | 0.34 | 0.20 | 1.66 | 42.29 | 0.1049 | 0.33 | 0.20 | 1.60 | 42.20 | 0.1167 |
| **Time (weeks)** | 0.00 | 0.00 | -0.75 | 137.48 | 0.4565 | 0.00 | 0.00 | -0.70 | 137.35 | 0.4832 |
| **Age** | -0.01 | 0.02 | -0.72 | 39.61 | 0.4782 | -0.01 | 0.02 | -0.65 | 39.63 | 0.5181 |
| **Motion** | 0.16 | 0.18 | 0.91 | 169.12 | 0.3629 | 0.13 | 0.17 | 0.77 | 167.97 | 0.4445 |

| | Distance (VWFA2-VWFA1) ~ Time (session) + Age + Motion + ( 1 \| sub | | | | | | | | | |
|---|---|---|---|---|---|---|---|---|---|---|
| | Cosine Distance | | | | | Pearson's distance | | | | |
| | β | se | t | df | p | β | se | t | df | p |
| **(Intercept)** | 0.34 | 0.20 | 1.68 | 42.57 | 0.1007 | 0.33 | 0.21 | 1.62 | 42.46 | 0.1128 |
| **session: post** | -0.01 | 0.02 | -0.38 | 135.64 | 0.7040 | -0.01 | 0.02 | -0.31 | 135.48 | 0.7568 |
| **session: 6mo** | 0.01 | 0.02 | 0.48 | 135.59 | 0.6326 | 0.01 | 0.02 | 0.55 | 135.45 | 0.5837 |
| **session: 12mo** | -0.03 | 0.02 | -1.19 | 135.13 | 0.2355 | -0.02 | 0.02 | -1.18 | 135.02 | 0.2392 |
| **Age** | -0.01 | 0.02 | -0.75 | 39.62 | 0.4589 | -0.01 | 0.02 | -0.68 | 39.63 | 0.4987 |
| **Motion** | 0.17 | 0.18 | 0.95 | 166.88 | 0.3434 | 0.14 | 0.17 | 0.81 | 165.70 | 0.4185 |

**Table S6.** Linear mixed effect models testing whether the distance between VWFA-2 and VWFA-1 in connectivity space changes over time in the intervention group. Age is years at the pre-intervention visit. Motion- mean framewise displacement (mm).
